# Novel ratio-metric features enable the identification of new driver genes across cancer types

**DOI:** 10.1101/2020.01.17.910075

**Authors:** Malvika Sudhakar, Raghunathan Rengaswamy, Karthik Raman

## Abstract

An emergent area of cancer genomics has been the identification of driver genes. Driver genes confer a selective growth advantage to the cell and push it towards tumorigenesis. Functionally, driver genes can be divided into two categories, tumour suppressor genes (TSGs) and oncogenes (OGs), which have distinct mutation type profiles. While several driver genes have been discovered, many remain undiscovered, especially those that are mutated at a low frequency across samples. The current methods are not sufficient to predict all driver genes because the underlying characteristics of these genes are not yet well understood. Thus, to predict novel genes, we need to define new features and models that are not biased and identify genes that might otherwise be overshadowed by mutation profiles of recurrent driver genes. In this study, we define new features and build a model to identify novel driver genes. We overcome overfitting and show that certain mutation types such as nonsense mutations are more important for classification. Some known cancer driver genes, which are predicted by the model as TSGs with high probability are ARID1A, TP53, and RB1. In addition to these known genes, potential driver genes predicted are CD36, ZNF750 and ARHGAP35 as TSGs and TAB3 as an oncogene. Overall, our approach surmounts the issue of low recall and bias towards genes with high mutation rates and predicts potential novel driver genes for further experimental screening.

## BACKGROUND

Cancer is one of the leading causes of morbidity globally, with more than 18.1 million cases reported in the year 2018 [1]. A major focus of cancer research has been the understanding of molecular mechanisms that govern tumorigenesis and the targets that can be used for treatment. Cancer cells are distinct because of their genomes, which give these cells the ability to divide and metastasize to other tissues in the body. It has been observed that mutations in some genes [2, 3] confer the ability of oncogenesis to these cells. The term “driver” was coined to refer to mutations in the genome that pushed the cell to oncogenesis [4]. Of all the mutations present in a cancer cell, not all are involved in giving a cellular advantage to the cell to divide uncontrollably. Driver mutations [4, 5] are those that were advantageous for tumour development and metastasis during the clonal evolution [6, 7]. On the other hand, *passenger* mutations [4, 5] are mutations that are accumulated during normal cell division or due to high mutational rates in cancer cells, but their presence or absence does not affect the progression and establishment of tumours.

Driver genes are effectively those genes that harbour mutations that provide them with a selective advantage to divide and grow unchecked. These driver genes not only help the cells bypass the cell cycle checkpoints to divide in an uncontrolled fashion but also give added functionality, such as bypassing the immune system [8, 9] and angiogenesis [10, 11], which lead to their persistence in the body. While certain cancers with well-understood mechanisms show that the presence of driver mutations is recurrent in most samples of a cancer type [2], others seem to have mutations that occur at a lower frequency. Driver genes that contain lower frequency of mutations are difficult to identify [12] because most likely these genes work in combination with other genes to confer a selective advantage to the cell.

Driver genes can be of two kinds depending on the role of the gene in a normal cell type. A tumour suppressor genes (TSG), as the name suggests, is the cell’s defence mechanism from becoming a cancer cell. When such a gene loses its function due to say, frameshift mutations or nonsense mutations, a selective growth advantage is conferred to the cell. Proto-oncogenes undergo gain of function mutations to become into an oncogene (OG). Mutations in both TSGs and OGs tip the balance of a normal cell into becoming a cancer cell. While many TSGs and OGs have been discovered for different cancer types, most of them are highly potent and recurring in different patients. A pan-cancer model will help in identifying patterns which might be lost while studying a cohort or specific cancer type, owing to low sample sizes or mutation frequencies. A key aim of this study is to find low-frequency driver genes by classifying them into TSGs and OGs.

There are broadly two classes of methods for identifying driver genes based on mutational data. The first class of methods [13–15] rely on the rate of mutations in genes for a set of patients, to identify driver genes. In these studies, the background mutation rate is estimated, and genes that show statistically different mutation rates are identified as driver genes. The rate of different types of mutations is used to calculate the background mutation rate [14, 15]. The methods of identification differ in the statistical method used [14]. The rate of cell division and length of the gene needs to be taken into account as the mutation rate may change depending on cell type and length and position of the genes [15].

Among the different methods that exist for identifying driver genes, when validated using the Cancer Gene Census (CGC) [16], it was observed that while the precision of identifying these genes was high, they had a very low recall [12]. Furthermore, genes identified through these approaches have a high <u>recurrence of being mutated</u> across different tumour samples. We now know that the rate of mutation is not sufficient for the identification of driver genes; instead, genes with low mutation rate can be driver genes if a mutation occurs at functionally important positions.

The second class of methods use a ratio-metric approach, where not only the repeated occurrence of mutations is taken into consideration, but also the functional impact of the mutations. Ratio-metric algorithms [17–19] capture the proportion at which the different mutation types occur. The type of mutations and their ratios vary and are distinct for TSGs and OGs. For instance, TSGs are more likely to have indels (insertions and deletions), more specifically frameshift mutations, that lead to loss of function of the protein. On the other hand, OGs tend to accumulate missense mutations that confer the protein with a “gain of function” [5, 20]. These features are then used for differentiating between these two types of driver genes.

While these methods do capture some mutation patterns observed across samples, low recall shows that our understanding of the characteristics that define TSGs and OGs is far from complete. In this study, we define new features that calculate entropy and frequency of different mutation types along with other ratio-metric features. Our aim is to identify important features for TSGs and OGs that can help classify a given gene as a TSG or an OG. Since the ratio-metric approach is based on the type of mutations and these differ for TSGs and OGs, genes were classified into two classes. Further, classification problems are prone to overfitting resulting in high classification scores in the training set, but the model can turn out to be unreliable for predictions using new data. We outline a method for estimating parameters for the given classification algorithm and avoid overfitting. We use the final model to predict novel driver genes by classifying a list of unlabelled genes; we validated our predictions by illustrating the presence of known TSGs and OGs among our predictions and through functional analysis of the predicted novel genes. We calculated the mutation rates and compared our results with the widely used tool MutSigCV and show that our method is able to pick out many driver genes that have very low mutation rates. Further, we used a pan-cancer model to predict driver genes that were tissue-specific.

## RESULTS

We define novel features and a method to estimate parameters and build a classifier using pan-cancer data to predict TSGs and OGs. The classifier is further used to predict labels for unlabelled genes, at pan-cancer and tissue-specific levels, which are analysed for functional enrichment.

### Novel features used for classification of TSGs and OGs

We trained multiple random forest models using a subset (80%) of 136 TSGs and 76 OGs for each fold of the cross-validation. We performed a five-fold cross-validation while estimating hyper-parameters for the model followed by multiple random iterations to estimate stable hyper-parameters and avoid overfitting (as defined in Methods). It is important to carefully consider overfitting as the initial training set is not very large. The accuracy for the test set reduces compared to the training set, but this difference is not substantial. We note that TSGs can be predicted with higher accuracy than OGs; it is probable that the features are biased at capturing information regarding TSGs better than OGs. Across the multiple models, an average accuracy of 0.76 ± 0.03 was achieved. These models were further used for the identification of novel genes as well as tissue-specific analyses. Our model presents a significant improvement in recall for TSGs. For OGs, the recall is similar to those observed in other tools. Nevertheless, an average recall of driver genes (comprising both classes) shows an improvement over the tools reported earlier [12].

To identify features important for the classification of TSGs and OGs, we calculated the average rank of each feature, across all models. We observe that the top-ranking features contain LOF and missense mutations (Supplementary Table S1). The new features that replace old features in the top 18 ranks are Nonsense entropy, High missense frequency, Compound/benign, High Frameshift Frequency, Damaging/kb, Compound/kB, Damaging/LoFI and HiFI/benign. Further, we used the training set genes to compare the distribution of feature values in TSG and OGs, and observed that our top-ranking features show the highest differences between the two distributions (Fig 1). While it is common knowledge that LOF mutations accumulate in TSG and recurrent missense mutations in OGs, we formally show that the feature distribution is different for these two functional classes.

**Figure 1.**
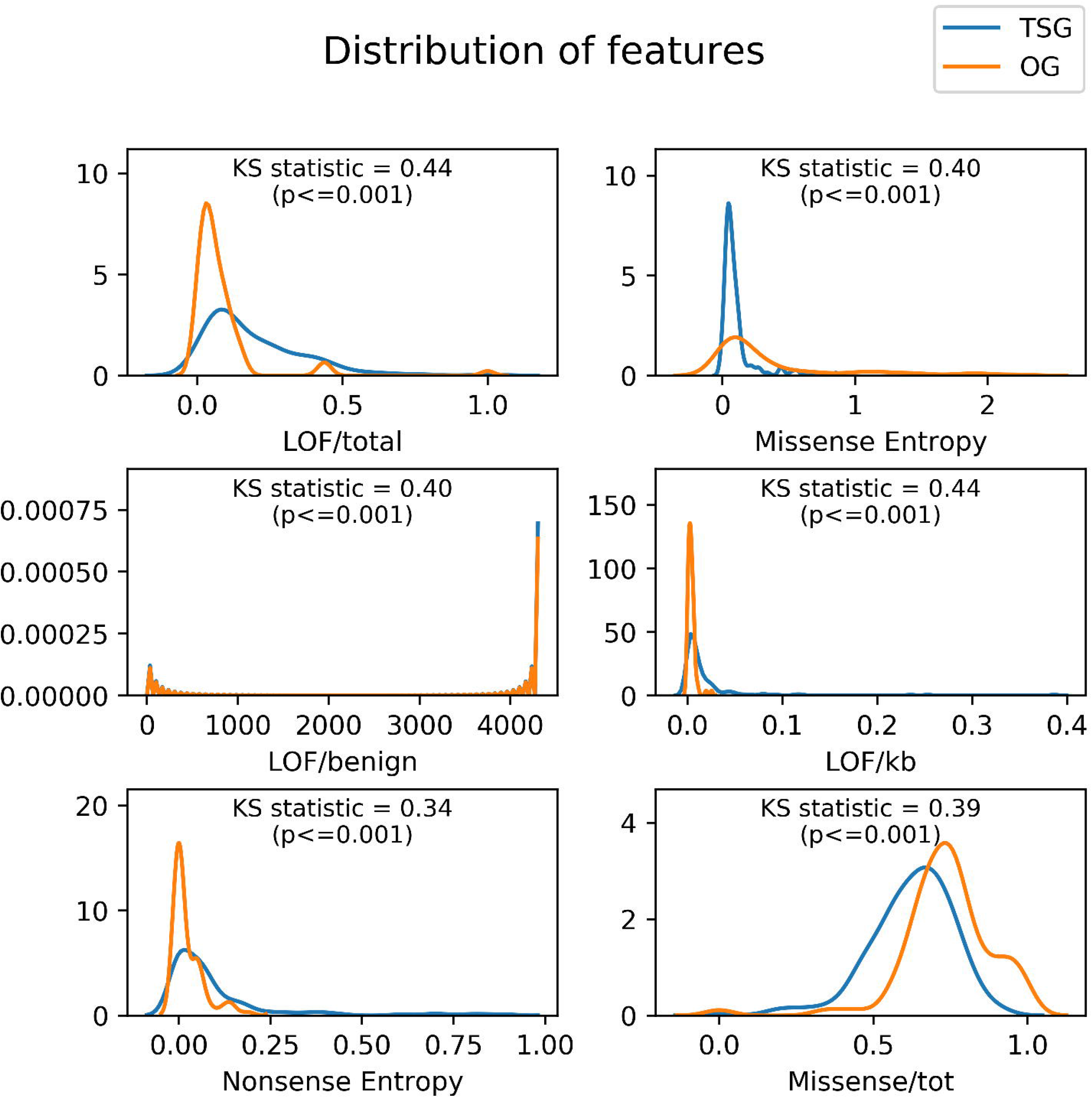
Distribution of top features identified by the classifier for TSG and OG. Training genes were used to study the differences between the distributions of features (kernel density) in TSG and OG. Kolmogorov-Smirnov statistic and the p-value is given for each feature. Higher value of KS statistic shows magnitude of difference of the two distributions.

### Iterative hyper-parameter estimation avoids overfitting

Initial analysis for a large number of *n_estimator* for random forest and using BalancedBagging to manage class imbalance gave higher accuracy score for training sets comparable to Davoli *et al.*, (2013). However, these showed very low accuracy for the test set (Table 2), indicating overfitting. Additionally, we observed that changing the random seed showed substantial variation in results. This variation is unexpected and could perhaps stem from non-optimum parameters used for classification or the small size of the data. To avoid this variation, we re-estimated the random forest parameters, *n_estimator*, *max_features*, *max_depth* and *criterion*. Changing the *n_estimator* had a major effect on classification, and grid search with cross-validation did not help in removing overfitting.

**Table 1.** Classification metrics for training and test set. Numbers in bold indicate best performances for each metric between TSG and OG. The metrics are standard, and are defined as follows (T stands for True, F for false, P for positives and N for negatives): Accuracy = (TP + TN)/(TP + FP + TN + FN); Precision = TP/(TP+FP); Recall = TP/(TP+FN); F1 score is the harmonic mean of Precision and Recall.

|  |  | Accuracy | F1 score | Precision | Recall |
| --- | --- | --- | --- | --- | --- |
| Training set | OG | 0.86 ± 0.04 | 0.77 ± 0.07 | **0.93 ± 0.04** | 0.67 ± 0.09 |
|  | TSG |  | **0.90 ± 0.03** | 0.84 ± 0.04 | **0.97 ± 0.01** |
| Test set | OG | 0.76 ± 0.03 | 0.59 ± 0.10 | **0.79 ± 0.12** | 0.50 ± 0.19 |
|  | TSG |  | **0.83 ± 0.02** | 0.77 ± 0.07 | **0.91 ± 0.07** |

**Table 2.** Classification metrics for training and test set using BalancedBagging.

|  |  | Accuracy | F1 score | Precision | Recall |
| --- | --- | --- | --- | --- | --- |
| Training set | OG | **0.93 ± 0.05** | 0.92 ± 0.06 | 0.86 ± 0.09 | 0.99 ± 0.01 |
|  | TSG |  | 0.94 ± 0.04 | 1.00 ± 0.01 | 0.90 ± 0.07 |
| Test set | OG | **0.69 ± 0.06** | 0.64 ± 0.06 | 0.56 ± 0.07 | 0.75 ± 0.06 |
|  | TSG |  | 0.73 ± 0.06 | 0.82 ± 0.04 | 0.65 ± 0.09 |

We overcame this by multiple iterations of hyper-parameter estimation by changing the random seed, which helps us identify more stable hyper-parameters. This gave lower accuracy for training sets but improved the accuracy of the test set considerably. When varying sets of random seeds (10, 20, 40, 80, 160, 320) were used, the results were consistent across all cross-validation folds (test set accuracy 0.76 and standard deviation 0.03) implying the increasing number of random seed iterations do not decrease or improve accuracy. We observe that for a given data fold, the hyper-parameters selected are more stable for varying sets of random seeds. While different parameter sets dominate as the data is changed, the overall results on the test set do not vary.

### Model identified novel TSGs and OGs along with known driver genes

All genes that were not used for training the models were classified into TSGs and OGs. This list also contained genes that are known driver genes present in CGC but not used for training. The labels were predicted for the unlabelled genes, of which 126 genes or transcripts showed consensus across all models. CGC known driver genes contributed to 40.5% of these predictions which included genes such as ARID1A, ATRX, NF1, TP53, RB1, and STAG1 and their transcripts. Some novel genes predicted consistently are SIN3A, ZNF750, IWS1, CD36, ARHGAP35, MGA, and RASA1 as TSGs. The model tends to be biased towards TSGs with 699 genes with consistent predictions for 3 or more models out of which only 9 are predicted as OGs. The top OGs predicted are U2AF1, BCL2L10, KRAS, MAP1LC3B, C11orf68, TAB3, MED12, MAX, and BRAF. Further, we show not all transcripts of a gene behave like a driver gene, for e.g. ATRX transcript ENST00000373344 is labelled as TSG but not ENST00000400866, ENST00000373341. The presence of known driver genes among top TSG and OG shows the validity of the model and those other genes in the list are potential driver genes.

Enrichment analysis of genes for various KEGG and BIOCARTA pathways revealed genes involved in different cancer pathways such as myeloid leukaemia, and pancreatic cancer. Genes are also enriched for various signalling pathways associated with cell growth, such as EGF and PDGF signalling pathways. Further, to validate, a similar analysis was conducted using genes used for training the model. We find GO terms related to cell cycle, regulation of transcription, signalling and cell cycle arrest to be common for both results. These keywords were further clustered with top clusters associated with genes involved in zinc-finger proteins, helicases, ATP-binding, ARID binding and cancer pathways. The analysis shows known driver genes and predicted driver genes enrich for similar pathways.

### Our approach identifies genes with low mutation frequency

We analysed the mutation frequencies of the predicted genes. Mutation rates were calculated using MutSigCV, a well-known driver gene predictor, which calculates mutation rates to identify driver genes. MutSigCV ranks all genes of which a total of 602 driver genes were identified above the threshold (p <=0.005, q <= 0.01). Training data labels were used to compare the two methods. MutSigCV identified 40% for our training gene set with 85 genes predicted as driver, while our model did better by predicting 85% of genes. The mutation rates of the genes predicted by the two models were compared. Since MutSigCV ranks all genes, we picked top genes equal in size to our model predictions (>=5 model consensus) and calculated KS statistic against training set and plotted the fraction of genes below mutation rate of each gene. We observe that distribution of mutation rates is similar to training set for our predicted genes, while MutSigCV tends to be biased towards genes with higher mutation rates (Fig 2). The minimum mutation rate predicted for our model was 0.35 while for MutSigCV was 0.90. The KS (Kolmogorov-Smirnov) statistic for both models when compared to training set shows the difference is far lesser for our model when compared to MutSigCV (Table 4), which shows that the distribution of mutation rates is similar to what is expected.

**Figure 2.**
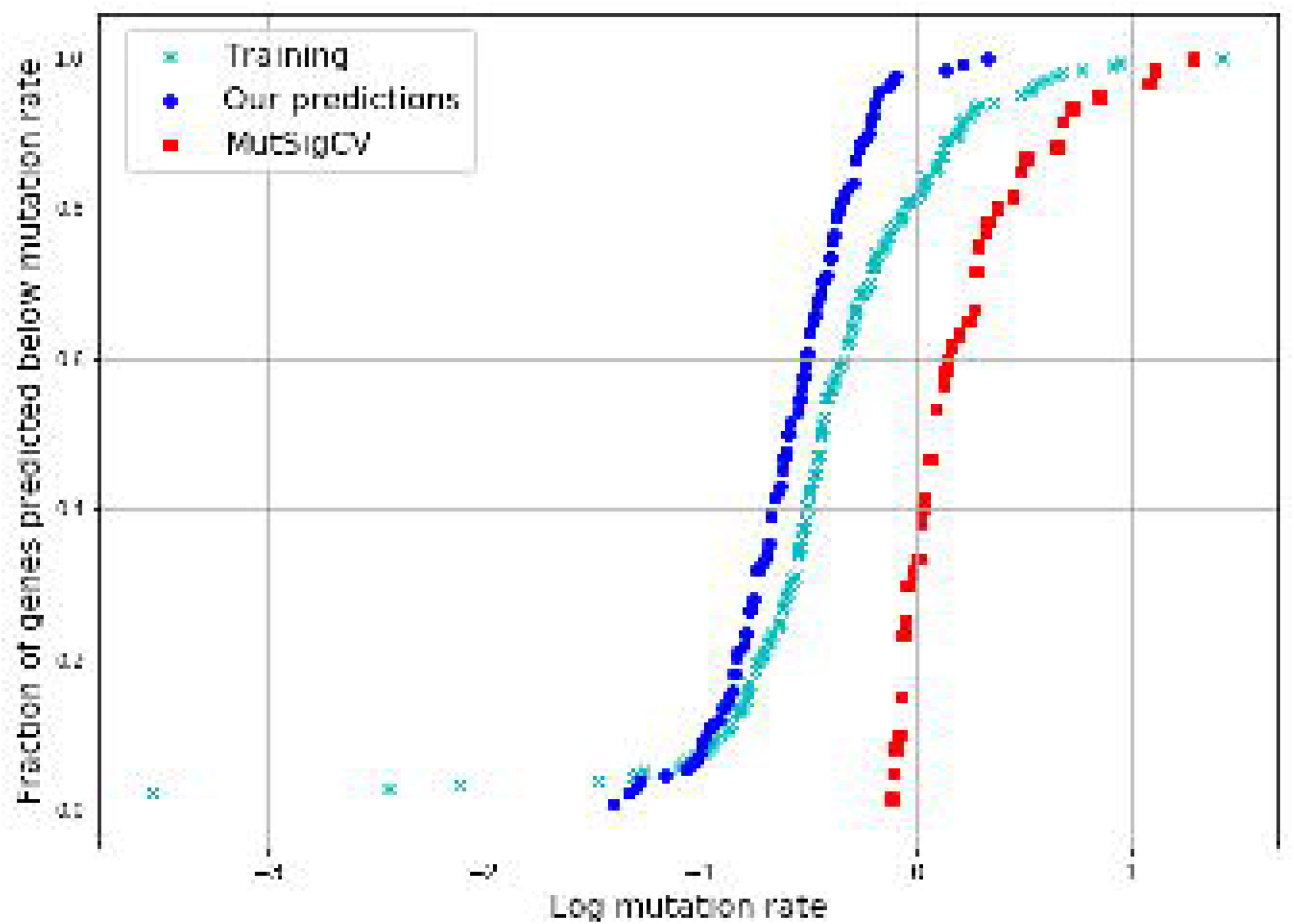
Fraction of genes predicted plotted against log transformed mutation rates. Genes predicted by a given method were sorted based on their mutation rate and plotted against the fraction of genes predicted below the given mutation rate

**Table 3.**
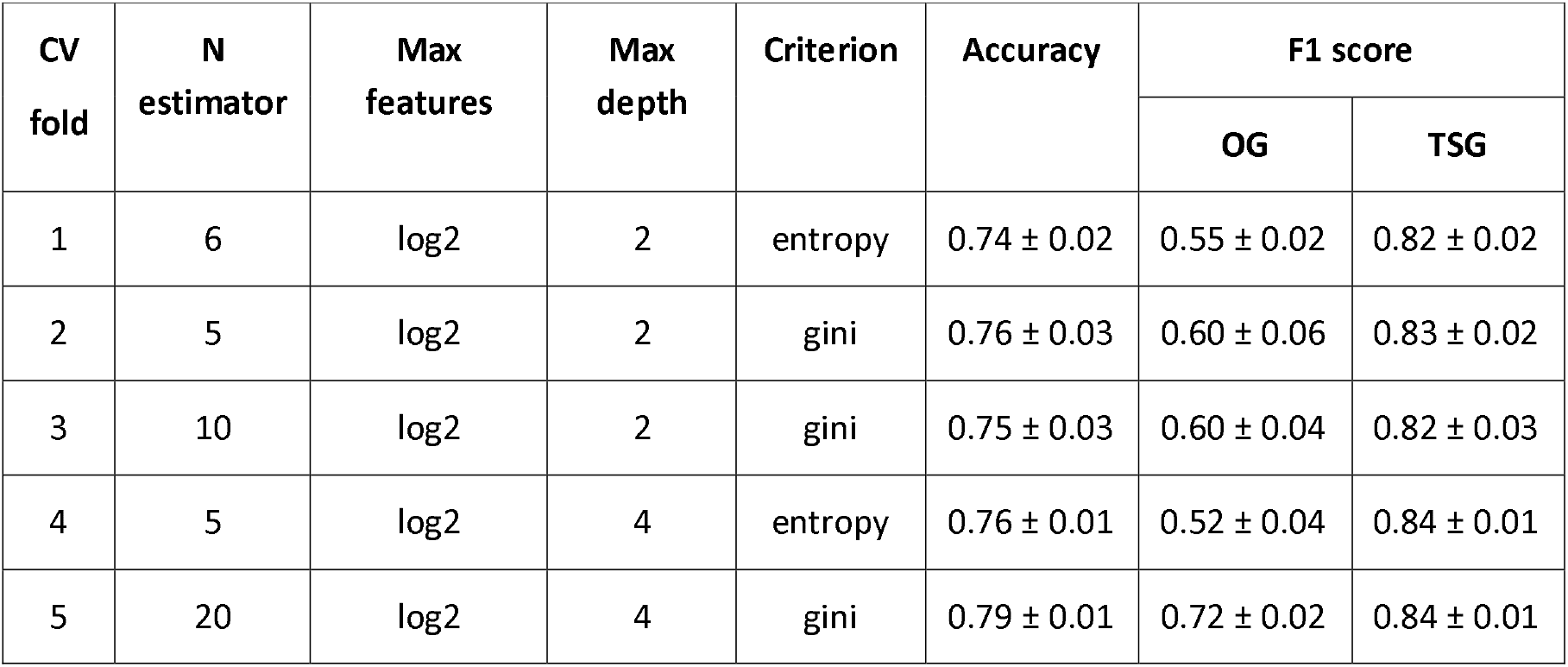
Hyper-parameters for each of the folds. For each cross-validation fold, the most frequent hyper-parameter set is reported. The average accuracy and F1-scores across the different random seed iterations (10, 20, 40, 80, 160, 320) are given along with the standard deviation.

| CV fold | N estimator | Max features | Max depth | Criterion | Accuracy | F1 score |  |
| --- | --- | --- | --- | --- | --- | --- | --- |
|  |  |  |  |  |  | OG | TSG |
| 1 | 6 | log2 | 2 | entropy | 0.74 ± 0.02 | 0.55 ± 0.02 | 0.82 ± 0.02 |
| 2 | 5 | log2 | 2 | gini | 0.76 ± 0.03 | 0.60 ± 0.06 | 0.83 ± 0.02 |
| 3 | 10 | log2 | 2 | gini | 0.75 ± 0.03 | 0.60 ± 0.04 | 0.82 ± 0.03 |
| 4 | 5 | log2 | 4 | entropy | 0.76 ± 0.01 | 0.52 ± 0.04 | 0.84 ± 0.01 |
| 5 | 20 | log2 | 4 | gini | 0.79 ± 0.01 | 0.72 ± 0.02 | 0.84 ± 0.01 |

**Table 4.** Kolmogorov-Smirnov statistic for mutation rate distribution of predicted genes when compared to training set genes. KS statistic for the top 60 predicted genes when compared with 208 genes in the training set.

| Method | KS statistic | p-value |
| --- | --- | --- |
| MutSigCV | 0.774 | <<0.001 |
| Our model | 0.193 | 0.054 |

### Driver genes are tissue-specific

Cohort studies tend to be specific to a cancer type. The usefulness of a pan-cancer model is further elucidated when it can be used to identify tissue-specific driver genes. The objective of predicting genes using a subset of data specific to tumour primary tissue source was to identify genes specific to a cancer type. This helped in identifying genes which might otherwise be lost in biological noise (Table 5). We observe TP53 predicted as TSG across the different tissues. Other known driver genes that weren’t identified by the pan-cancer analysis were identified such as CBFB, CDH1, PTEN in breast cancer and APOB in liver. Genes such FAM182A, SOX9, AHNAK2, ENSG00000121031, FLT3LG, PMEPA1, ZFP36L2 in the large intestine, ALB, KRTAP19-1, APOB, CD200, CRYGD, KRTAP24-1, OR6N2 in the liver are novel predictions, and their functions in these cancers can further be studied. We used the pan-cancer models to predict tissue-specific driver genes and identified new genes not reported by the pan-cancer analysis.

**Table 5.** Driver genes predicted for each of the cancer types. The genes reported showed consensus for >4 CV models. Genes in bold did not find similar consensus in the pan-cancer predictions. Novel genes are underlined.

| Primary Tissue | Genes |
| --- | --- |
| Breast cancer | TP53, **CBFB**, RUNX1, **CDH1**, GATA3, **PTEN**, TBX3 |
| Central nervous system | TP53, **HCN1** |
| Endometrium | **KRAS**, PIK3R1, **PTEN** |
| Hematopoietic | TP53, B2M, **CCND3**, HLA-A |
| Kidney | PBRM1, **VHL**, TP53 |
| Large intestine | TP53, FBXW7, <u>**FAM182A**</u>, <u>SOX9</u>, <u>**AHNAK2**</u>, TCF7L2, <u>ENSG00000121031</u>, <u>**FLT3LG**</u>, <u>**PMEPA1**</u>, <u>**ZFP36L2**</u> |
| Liver | TP53, <u>**ALB**</u>, <u>**KRTAP19-1**</u>, <u>**APOB**</u>, <u>**CD200**</u>, <u>**CRYGD**</u>, <u>**KRTAP24-1**</u>, <u>**OR6N2**</u> |

Genes identified for breast cancer was validated by supporting literature. CBFB [21] and PTEN [22, 23] is a known TSG in breast cancer. PTEN is found to be under-expressed in breast cancer [24, 25]. While CDH1 mutations are found mostly in stomach cancer, they are also shown to be frequently occurring in lobular breast cancer [26, 27]. Pathway analysis of breast cancer genes shows enrichment of pathways involved in gene expression regulation governed by TP53, RUNX1 and PTEN which includes pathways that regulates estrogen-mediated transcription. CBFB deletion leads to expression loss of RUNX1[21], which can no longer regulate NOTCH signalling by repression, which is confirmed by pathway analysis. Some apoptosis pathways are enriched that include CDH1 and TP53 genes. The genes identified by the pan-cancer model for breast cancer samples identify genes functionally important in breast tumour cells.

Predictions made for liver cancer were mostly novel, which made literature validation difficult. RNA expression levels of genes APOB, ALB and CD200 were higher compared to all other tissues (as reported by The Human Protein Atlas). Higher albumin levels are known to decrease the risk of HCC (Hepatocellular carcinoma) [28]. APOB mutational signatures are shown computationally to be significant to predict prognosis, by loss of regulation of genes such as TP53, PTEN, HGF [29]. While role of other genes is difficult to elucidate, our method helps identify research gaps which can be filled by studying these potential driver genes.

## DISCUSSION

Identification of driver genes has been an important focus area of cancer research because these genes are potential targets for therapy and biomarkers. Different approaches have been used for identification using mutational information [17, 18, 30], gene expression levels [31], protein structural information [32], network analysis [33, 34] or using multiple data sources [31]. Advances in sequencing technologies have made mutational information easily available, and different tools have been developed to identify driver genes. Driver genes are further classified into TSGs and OGs based on the functional impact of the mutations they harbour.

We adopt a classification approach that is able to predict TSGs and OGs by leveraging a set of ratio-metric and other new features. Traditional methods identify genes based on the mutation rate. Compared to previous approaches, we ascribe a higher significance to functional impact along with the position of the mutations, as the genes might contain mutations in functionally important regions even though the mutation rate may not be very different from the background mutation rate. Features like nonsense entropy, frameshift frequency captures the recurrence of a mutation when multiple samples are considered, thus taking into account the position at which the mutation occurs.

For classification, many different algorithms are available, but the performance of the algorithm is dependent on the data and estimation of parameters. It is especially important while solving biological problems, where the training data might be small, to build robust models. We tried the classification of genes using support vector machines (SVM), logistic regression, balanced bagging as well as random forest and found that random forest performed better in this case. Further, high performance on a given data might also be due to overfitting. We sought to avoid overfitting by performing a standard 5-fold cross-validation while estimating random forest parameters as well as multiple iterations for estimation of stable parameters. We developed a procedure to verify that the predictions are reasonably stable. An ensemble of models is used to make final predictions.

It is important that the estimated parameters are robust to changes in data. For random forest, we estimated four parameters out of which *n_estimator* seemed to have a large effect on the classification. For large values of *n_estimator*, we were able to show high accuracy scores similar to Davoli et al., (2013) but the accuracy scores for test set were much lower. We were not able to compare our performance on the test set with that of Davoli *et al* (2013), as their test set results have not been published. To build a better model that is not biased to data, we needed a more robust classifier, that is sufficiently generalized and not dependent on the training data.

The models generated were used to find which of the new features are important for classification. To evaluate the model, we used 5-fold cross-validation with 20% test dataset while maintaining the ratio between TSGs and OGs and calculated metrics such as accuracy and F1 score. Instead of AUROC (Area under Receiver Operating Characteristic), we chose to show accuracy and F1 score, as AUROC only helps in estimating if the model can separate the given classes but tells us very little about the classification power for each of these classes. The F1 score is calculated for each of the given classes and helps understand if the model is biased towards any one of the classes. The accuracy score on the test set shows that mere accuracy is not sufficient for judging a model. The models perform slightly better for TSGs, though it is far poorer at classifying OGs.

While assessing the model, it is important to use metrics such as F1 score, as it scores predictions for each of the classes. Studies reporting only AUROC statistics present an incomplete picture and are not effective in estimating the performance of the model, especially in datasets having a class imbalance [35]. This is evident when we compare AUROC of Balanced bagging model (0.76 ± 0.07) with our model (0.54 ± 0.07). AUROC gives measures the models ability to separate the classes and not the prediction power. By reporting both accuracy as well as F1 score, we show that the model does not perform equally for both classes but tends to be better at classifying TSG than OG. This indicates that the chosen features are not sufficient to classify oncogenes.

Feature ranking shows that features containing information about LOF, nonsense, frameshift and missense mutations are important. Nonsense and frameshift mutations are frequently seen in TSGs while recurrent missense mutations are characteristic of OGs as they lead to “gain of function”.

The list of genes classified contained known driver genes and other transcript data for genes present in training and test set. We found that TSGs such as ATRX, PTCH1, and STAG2 were classified as TSGs with high probability. KDM6A gene and its transcripts (ENST00000377967, ENST00000382899) feature among the top, which shows that the model can also help classify a particular transcript of a gene. Similarly, TP53 and its six transcripts were all classified as TSGs. Genes U2AF1, KRAS, BRAF, MED12 and MAX were classified as OGs among the top genes identified as OGs. As the probability scores for OGs tend to be lesser than TSGs, relatively fewer OGs make the cut-off for the top 5 percentile.

Among the top TSGs identified, CD36 (previously known as FAT) is receptor protein for fatty acids. CD36 is also a prognostic marker for different cancer types [36, 37] and found in metastatic cells [36, 38]. While the expression of a gene is markedly different from normal cells, the molecular mechanism that enables metastasis is not well understood. Another gene, ARHGAP35, is a glucocorticoid receptor DNA binding factor, which has also been previously identified as a potential driver gene by other methods [39, 40]. ZNF750, zinc finger protein 750 has been established as a tumour suppressor in oesophageal squamous cell carcinoma [41–43] though it is absent from the CGC diver gene list. Some other potential TSGs not present in the CGC list are MBD6 and RASA1. In the human protein atlas, MAP1LC3B is labelled as a prognostic marker for renal and stomach cancer among the three shortlisted OGs.

Our model does have some limitations. We have used binary classification for identification of TSGs and OGs which, classifies all genes as either TSG or OG. All genes containing mutations are not driver genes, and thus, a majority of genes are neutral. We overcome this by taking consensus across the five models built. It may be possible to improve on this classification by solving a multi-class problem where each gene is identified as TSG, an OG or neutral gene. The difficulty in this problem stems from the huge class imbalance in the data as well as the definition of neutral genes. While there are studies showing the importance of a gene in tumour evolution, it is difficult to define genes that are not involved in cancer progression. Most methods use a list of genes that do not contain cancer driver genes and genes involved in cancer pathways, but this does not exclude potential driver genes.

Additionally, it has been seen that mutations are not always the reason for the change in functionality and regulation might also lead to change in expression at transcriptomic and proteomic levels. Other than adding new features to the analysis, including transcriptomic and proteomic data along with genomic mutation data might further improve the classification of genes.

## CONCLUSION

In summary, we see two main contributions of our paper. First, we developed a classifier, which enabled an improved recall of TSGs and OGs compared to previously proposed methods in the literature. We carefully avoided overfitting for achieving consistent and high confidence results. Second, we predicted many potential TSGs and OGs at both the pan-cancer and tissue-specific level, which form a ready short-list for further experimental investigation. Some of the top predictions were already well-known cancer drivers while others are reported in multiple cancer studies though their role in tumorigenesis is not yet well understood. Our approach is also readily amenable to the integration of other omic datasets, as they become available.

## METHODS

### Data

We downloaded somatic mutation data from Catalogue of Somatic Mutations in Cancer (COSMIC) (v79) [44]. These data were pre-processed to exclude hyper-mutated samples (samples containing more than 2000 mutations) Known SNPs were retained only if they were “confirmed somatic mutations”. The final processed data consist of 2,145,044 mutations from 20,667 samples across 37 primary tissues. COSMIC also contains transcript information, where different transcripts of a gene are saved as “gene_transcript” and are handled as separate genes. Splice site mutations were identified as mutations at 1 or 2 bps after the end of the exon border or 1 or 2 bps before the start of exon border. We used the popular tool Polyphen2 [45] to predict the phenotypic impact of missense mutations. For some mutations, Polyphen2 returns multiple scores, which we averaged for the purpose of our analyses.

TSGs and OGs for training and test were taken from the CGC [16] gene list. Only those genes that were labelled “TSG” or “OG” and not “Fusion” were used for this analysis. A total of 213 driver genes were used, of which 136 were TSGs and 77 were OGs. The TSG:OG ratio was maintained during all cross-validation steps and in both training and test sets.

### Ratio-metric features

Mutations were divided into 11 different categories [17, 45]: silent, missense, splicing, High Functional Impact (HiFI), Mid Functional Impact (MiFI), Low Functional Impact (LoFI), nonsense, frameshift, in-frame, nonstop or complex. Not all missense mutations are equally deleterious — labelling them into HiFI, MiFI and LoFI categories helps differentiate genes that have a large number of mutations with low impact, from genes that have relatively fewer mutations but with larger functional impact. We use PolyPhen2 scores to categorise mutations as HiFI (≥ 0.85), LoFI (≤ 0.15) and MiFI (between 0.15 and 0.85), to differentiate between high confidence pathogenic mutation predictions.

Additionally, other mutation categories were defined, which clubbed multiple mutations into one, such as ‘compound’ and ‘damaging’. Compound mutations are included because mutations types such as in-frame, nonsense and complex occur at a lower frequency than single nucleotide missense mutations, which might lead to patterns and impact of these mutations to be masked. Since the functional impact is similar to missense mutations, combining similar mutation types might help in capturing information of these less frequently observed mutation types. Loss of function (LOF) mutations introduce large changes into proteins, causing disruption of function. Damaging mutations are the sum of HiFI and MiFI mutations; these capture impact of multiple MiFI and sparse HiFI mutations. Many features compute a ratio of mutation types, as outlined in Table 6. We defined 37 features in all, with 18 of them being similar to those defined as Davoli *et al.*, (2013).

**Table 6.** Definitions of mutation categories and the ratio of mutation categories.

|  |
| --- |
| Compound mutations = missense + complex + inframe + nonstop – LoFI |
| Loss of Function (LOF) = nonsense + frameshift |
| Damaging = HiFI + MiFI |
| Benign = silent + LoFI |
| $ratio(A/B)_g = \begin{cases} \frac{A_g}{B_g} & \text{if } B_g \neq 0 \\ 2 * \max(A) & \text{if } B_g = 0 \end{cases}$ |

### Entropy and Frequency features

Entropy and frequency features were defined for four mutation types. A mutation (M_i_) in a given gene *i* is represented by its location. For missense mutations, M_i_ is represented as a tuple (*loc*, *wt*, *mt*) where *loc* is the location of the mutation, *wt* is the wild type nucleotide, and *mt* is the mutated nucleotide. If *k* unique mutations are present in a gene, *f_i_* gives the frequency for each of the mutations.

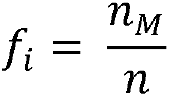

where *n_M_* is the number of occurrences of mutation *M* and *n* is the number of mutations in gene *i*.

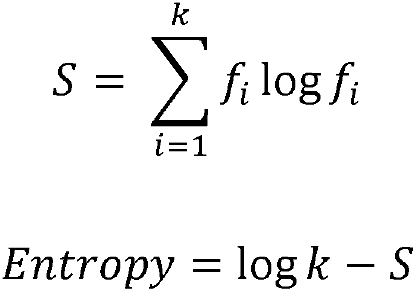

### Classification of genes

Different machine learning algorithms such as random forest, support vector machines and logistic regression were used, among which random forest gave the highest accuracy. Random forest was used for building a robust model and classifying TSGs and OGs. We used five-fold cross-validation to split data into training to test set ratio of 8:2; where each fold acts as a test set. We used the implementation of Random forest from the Python package Sci-Kit Learn [46]. We tuned the parameters using a five-fold cross-validation grid search along with multiple random iterations of random seed (described later). The parameters tuned are *n_estimator* (from 5-40), *max_features* (‘sqrt’ or ‘log2’), *max_depth* (2-4) and *criterion* (‘gini’ or ‘entropy’). The number of maximum features each decision tree considers is given by the parameter *max_features*, which can be calculated in two ways, as either the square root or log_2_ of the total number of features.

### Tuning hyperparameters and estimating the robustness of the classifier

Our initial results showed variation in classification depending on the random seed that was selected for classifying, even though cross-validation was used while estimating parameters. We used balanced bagging classifier to take into consideration the class imbalance and estimated parameters using cross-validation, which is the standard method. Poor results for this model led us to estimate hyper-parameters differently.

To avoid this variation, classification and parameter selection were done for multiple random seeds (Fig. 3 block B). Grid search with five-fold cross-validation was done for multiple different random seeds. Optimum parameters were selected by first estimating parameter *‘n_estimator’* and using it to estimate other parameters. Recurrence of ‘n_estimator’ across different random seeds was counted, and the maximum count was considered as the best ‘n_estimator’ to be given to the model. If multiple estimators were chosen, maximum accuracy during cross-validation was used to select one estimator. Maximum accuracy was used to find other parameters for the given best *‘n_estimator’*. The classification was rerun using the given parameters, and features were ranked. The model was used to predict the classification of test set genes.

**Figure 3.**
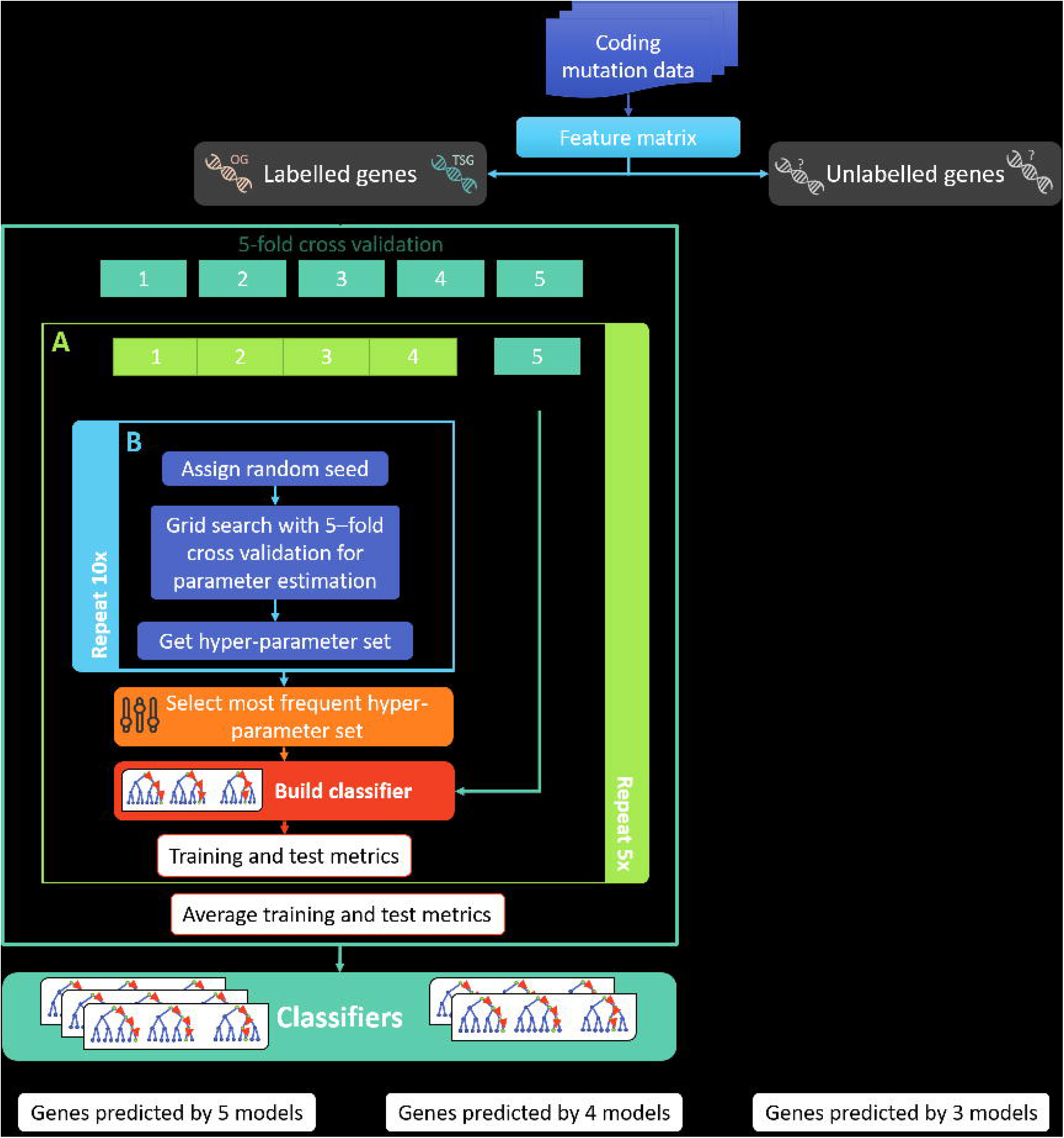
Methodology for identifying novel driver genes. The figure presents an overview of the different steps involved in our study. Block A (light green frame) shows how our classifier is built and is repeated 5 times. Block B (light blue frame) shows random iterations for estimation of hyper-parameters and is repeated 10 times.

To estimate the effect of the number of random iterations on parameter estimation, the classifier was built on a varying number of iterations of random seeds (10, 20, 40, 80, 160, 320). The stability of hyper-parameters selected was analysed based on the variation in the accuracy of the test dataset.

### Feature comparison and ranking

All features defined were used for classification and ranked depending on their contribution to the model. Average rank was calculated across the five validation sets. The features are given in Table 7.

**Table 7.** *The ratio-metric* features used in this study for classification.

|  |  |
| --- | --- |
| **Previously defined in the literature**<br>**(18 features)** | Silent/kb, Total Missense, Total Splicing, Total LOF, Missense/kb, LOF/kb, LOF/Silent, Splicing/Silent, Missense/Silent, LOF/Benign, Splicing/Benign, Missense/Benign, average Polyphen2 score, LOF/Total, Missense/Total, Splicing/Total, LOF/Missense, Missense entropy |
| **Defined in this paper**<br>**(19 features)** | HiFI/LoFI, HiFI/Benign, MiFI/kb, Nonstop/kb, Inframe/kb, Complex/kb, Compound/Benign, Compound/kb, Damaging/kb, Damaging/Benign, Damaging/LoFI, High Missense frequency, Frameshift entropy, High Frameshift frequency, Splicing entropy, High Splicing frequency, Nonsense entropy, High Nonsense frequency, Total MiFI |

### Identification and functional analysis of novel TSGs and OGs

We used the model built on the combined set of 37 features to classify unlabelled genes into TSGs and OGs. In total, 26,866 genes were classified as TSGs or OGs and ranked using their probabilities for each class. The genes given for classification contains different transcripts of the same gene symbol as different genes. In all, the gene list contained 18,951 unique gene symbols. Genes were labelled TSG and OG depending on their presence in the top 5 percentile and consensus across models built during cross-validation. Since not all genes are necessarily TSGs or OGs, genes which didn’t fulfil these criteria remained unlabelled. Novel TSG and OG gene list predicted by greater than four models were further used for functional analysis to find the major pathways and gene ontologies these genes are enriched for. Functional analysis was carried out using DAVID [47, 48] for both, the genes above the threshold as well as training set genes, and the results were compared.

Further, the pan-cancer classifier was used to predict genes in different cancer types based on the primary tissue where the tumour is formed. The data were filtered based on primary tissue, and the feature matrix was generated for tissues with >1000 samples. The data was then standardized and run using pan-cancer models described earlier.

We compared and calculated mutation rates using MutSigCV. Since the ground truth is not known for these predicted genes, we compared the genes used for training and calculated recall of these genes. Since MutSigCV does not classify genes as TSG or OG, the classes considered were Driver and Passenger. Further, we were interested in looking at the mutation rate distribution across the genes predicted. Since the distribution of mutation rates is unknown, we compared the similarity of the distribution of the predicted genes with the genes used for training (Kolmogorov-Smirnov statistic). Similarly, the similarity was compared for genes predicted by MutSigCV.

## LIST OF ABBREVIATIONS

AUROC: Area under Receiver Operating Characteristic
CGC: Cancer Gene Census
COSMIC: Catalogue of Somatic Mutations in Cancer
GO: Gene ontology
HCC: Hepatocellular carcinoma
HiFI: High Functional Impact
Indels: insertions and deletions
KS statistic: Kolmogorov-Simrnov statistic
LOF: Loss of function
LoFI: Low Functional Impact
MiFI: Mid Functional Impact
OG: oncogenes
TSG: tumour suppressor gene

## DECLARATIONS

### Ethics approval and consent to participate

Not applicable

### Consent for publication

Not applicable

### Availability of data and materials

Data for this analysis was downloaded from COSMIC (v79)

The processed data and codes are available in GitHub. (https://github.com/RamanLab/IdentifyTSGOG)

### Authors’ contributions

MS, RR and KR conceived and designed the study. MS, RR, and KR were involved in the analysis and interpretation of data. MS, RR and KR drafted the manuscript. The study was supervised by RR and KR. All authors read and approved the final manuscript.

## Acknowledgements

Not applicable

## ADDITIONAL FILE INFORMATION

Additional file 1: Table S1.

List of features and their ranks for each of the models and the calculated average rank. (XLSX 10.8 kB)

Additional file 2: Figure S1.

Distribution of features across the two classes for all the other features not included in Figure 1. (PDF 1.39 MB)

