## Supplementary material for "Novel ratio-metric features enable the identification of new driver genes across cancer types": Figure S1

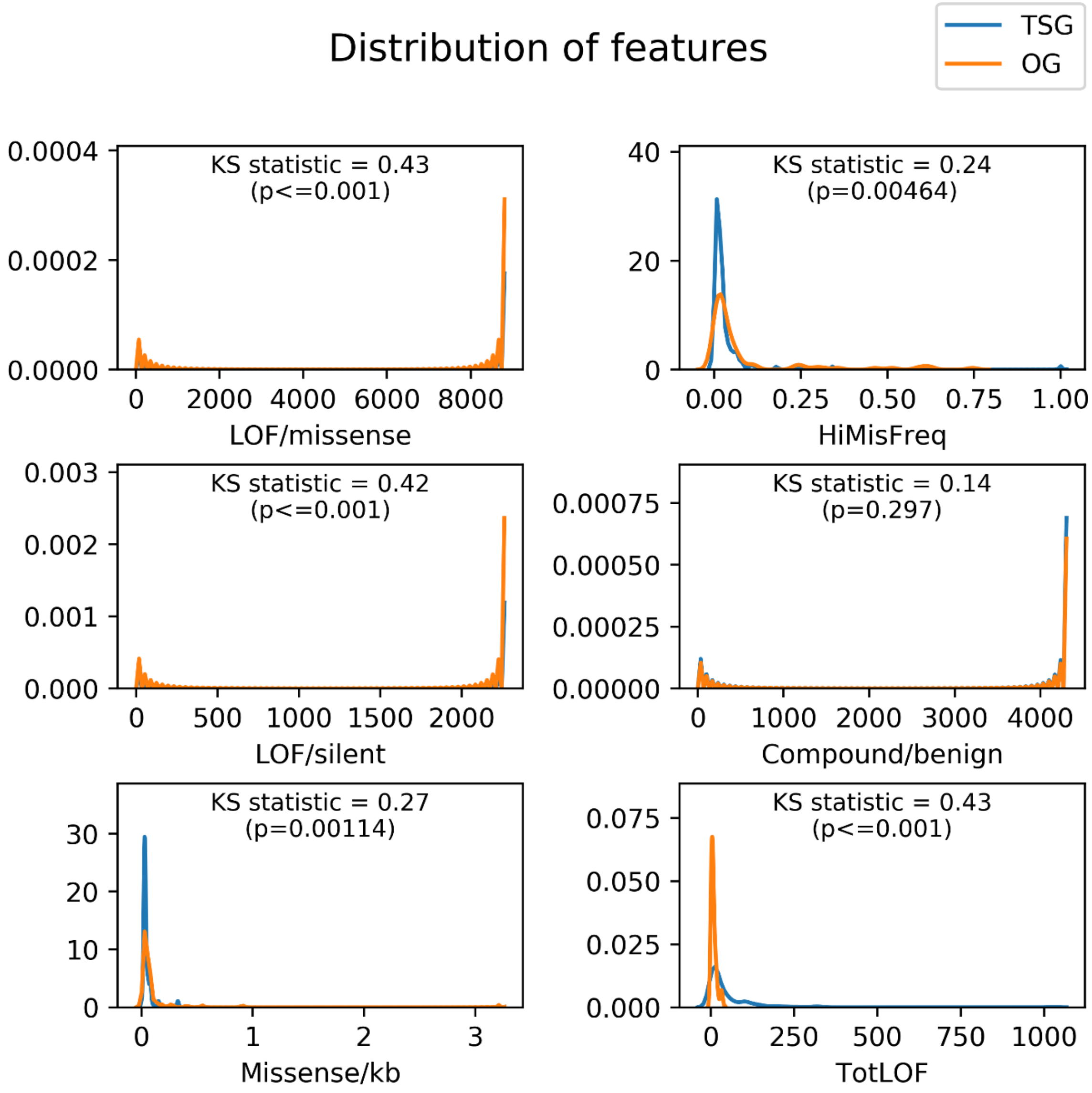


***Supplementary Figure 1a* Distribution of features used by the classifier for TSG and OG.** The kernel density was plotted for training genes used for classifying TSG and OG to study the differences between the distributions of features. Kolmogorov-Smirnov statistic and the p-value is given for each feature. Larger KS statistic means a larger difference in the two distributions.

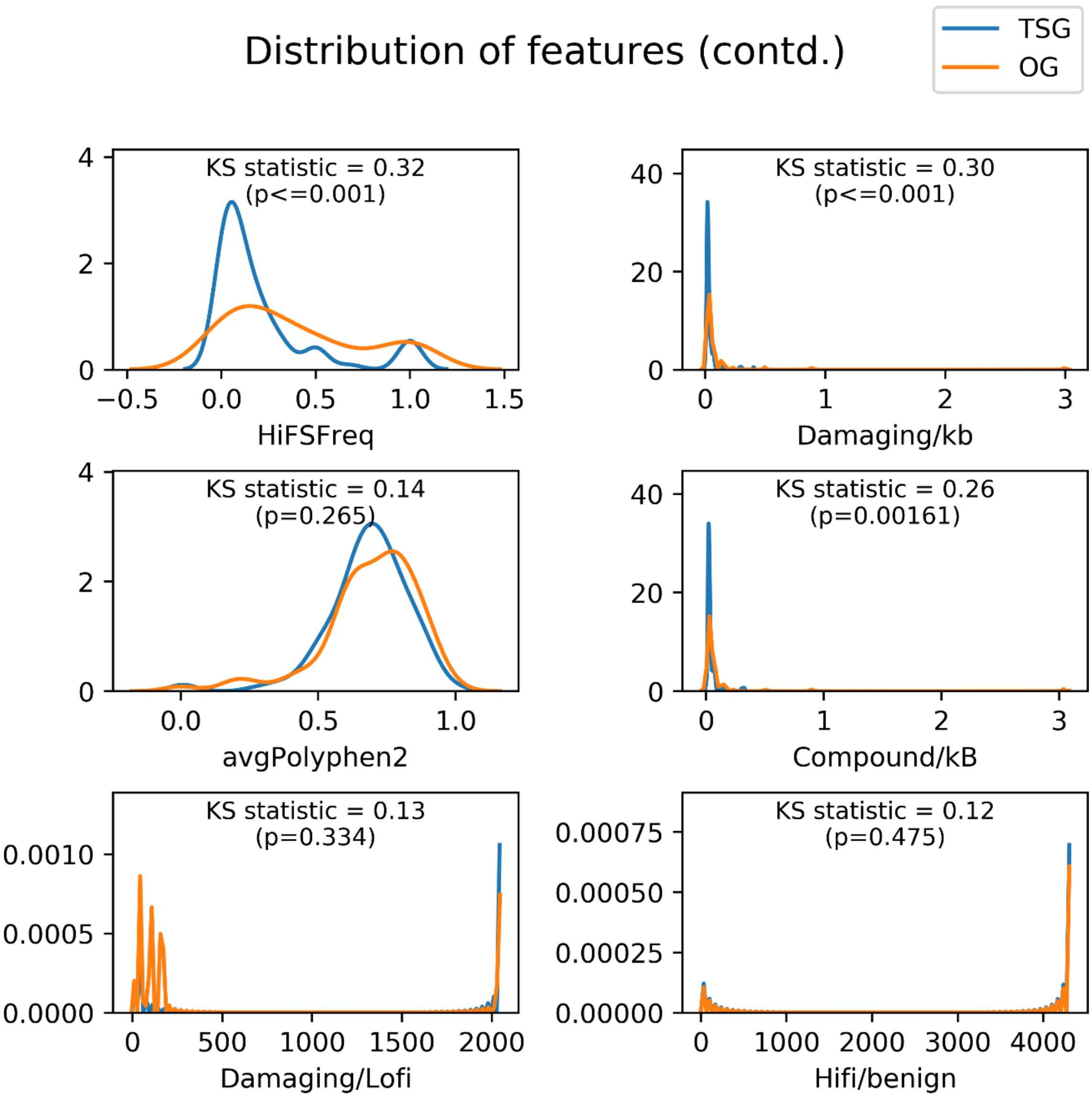


***Supplementary Figure 1b*** **Distribution of features used by the classifier for TSG and OG.**

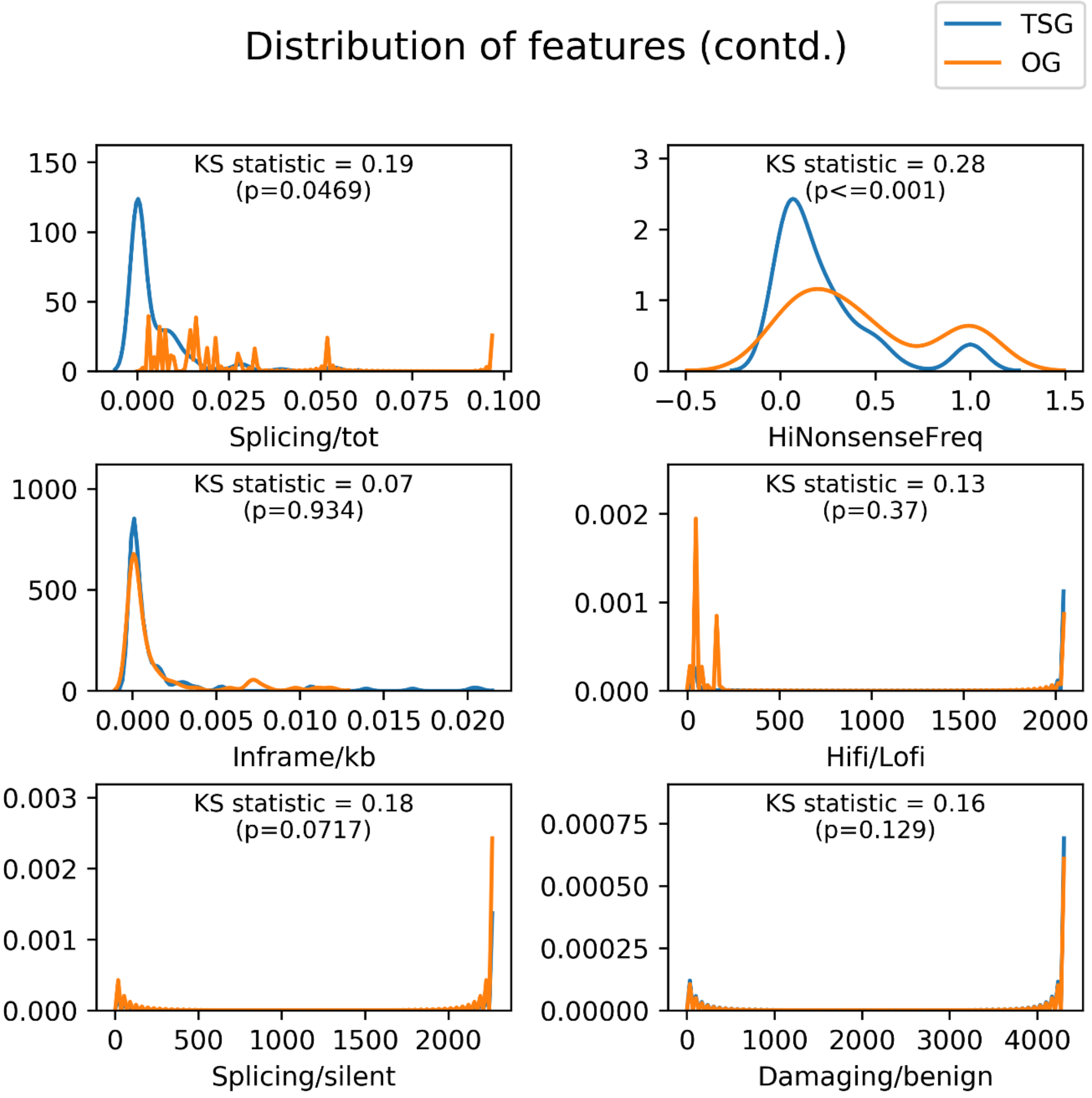


***Supplementary Figure 1c*** **Distribution of features used by the classifier for TSG and OG.**

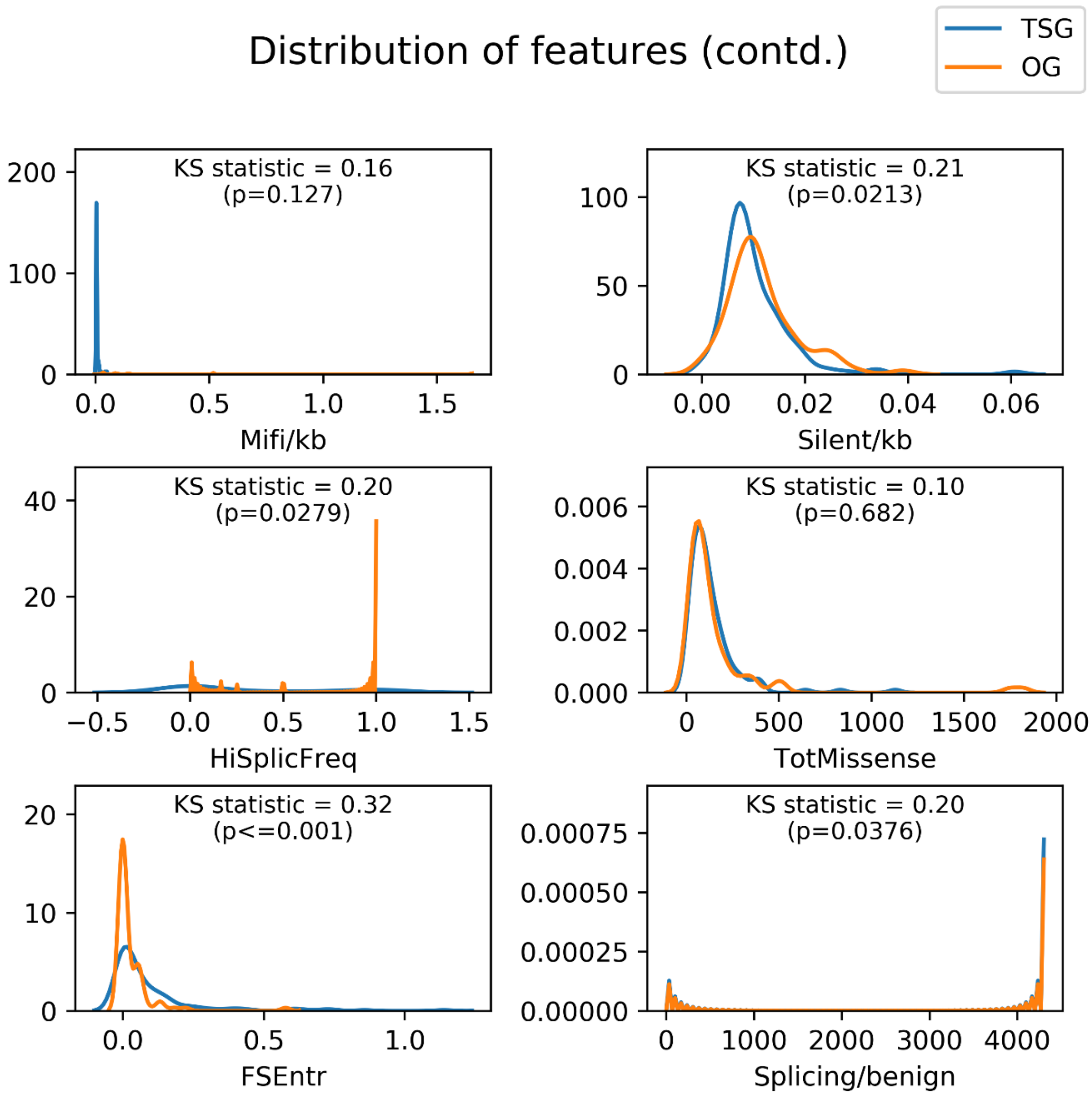


***Supplementary Figure 1d*** **Distribution of features used by the classifier for TSG and OG.**

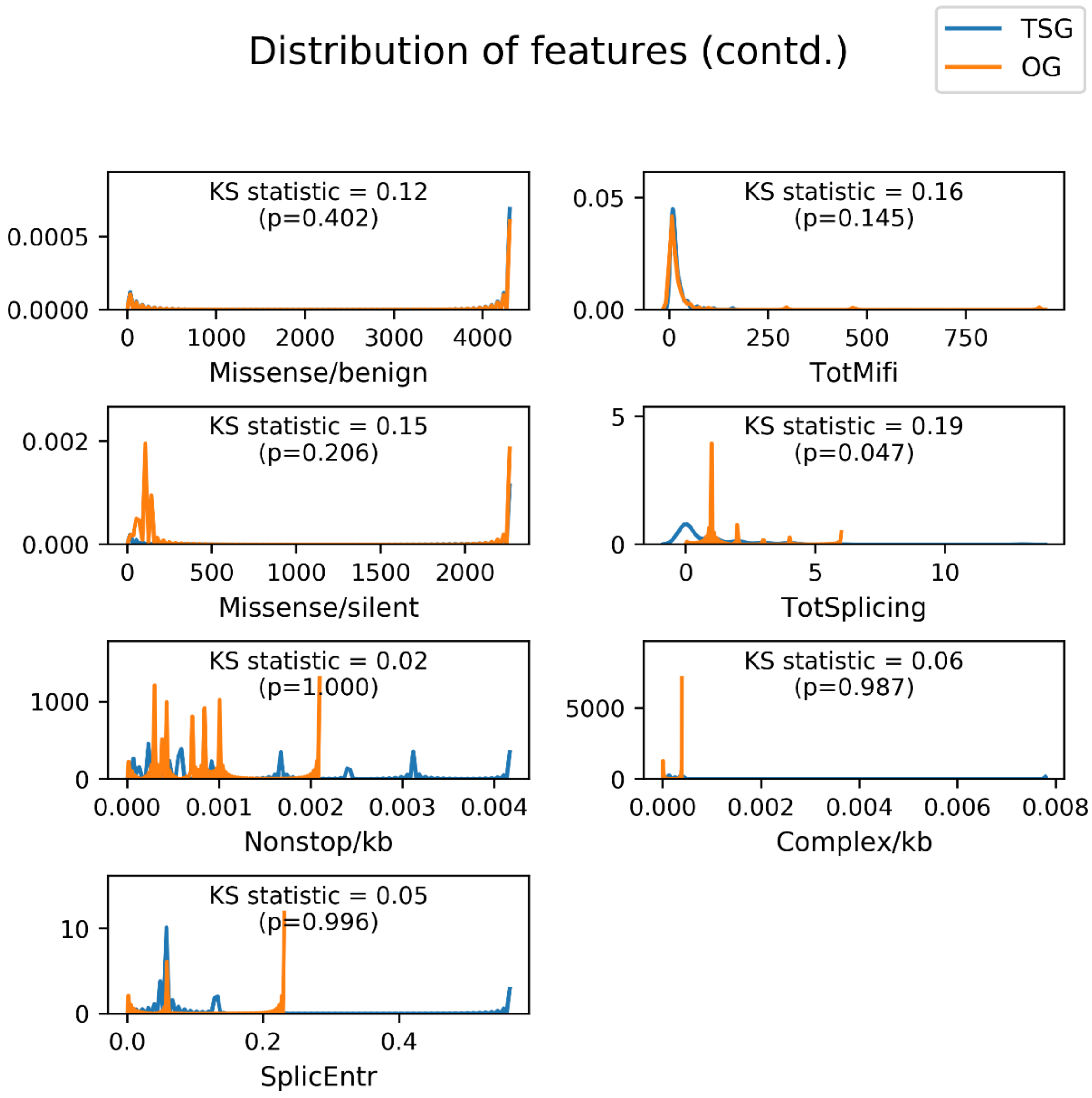


***Supplementary Figure 1e*** **Distribution of features used by the classifier for TSG and OG.**
